# EWS-FLI1 regulates and cooperates with core regulatory circuitry in Ewing sarcoma

**DOI:** 10.1101/2020.01.15.908475

**Authors:** Xianping Shi, Yueyuan Zheng, Liling Jiang, Bo Zhou, Wei Yang, Liyan Li, Lingwen Ding, Moli Huang, Sigal Gery, De-Chen Lin, H. Phillip Koeffler

**Affiliations:** Guangzhou Municipal and Guangdong Provincial Key Laboratory of Protein Modification and Degradation; State Key Laboratory of Respiratory Disease; Affiliated Cancer Hospital of Guangzhou Medical University; School of Basic Medical Sciences, Guangzhou Medical University, Guangzhou 510120, P.R. China; Department of Medicine, Cedars-Sinai Medical Center, Los Angeles, CA 90048, USA; Departments of Surgery and Biomedical Sciences, Cedars-Sinai Medical Center, Los Angeles, CA 90048, USA; Cancer Science Institute of Singapore, National University of Singapore, Singapore; School of Biology and Basic Medical Sciences, Soochow University, Suzhou 215123, P.R. China; National University Cancer Institute, National University Hospital Singapore, Singapore

## Abstract

Core regulatory circuitry (CRC)-dependent transcriptional network is critical for developmental tumors in children and young adults carrying few gene mutations. However, whether and how CRC contributes to transcription regulation in Ewing sarcoma is unknown. Here, we identify and functionally validate a CRC “trio” constituted by three transcription factors (TFs): KLF15, TCF4 and NKX2-2, in Ewing sarcoma cells. Epigenomic analyses demonstrate that EWS-FLI1, the primary fusion driver for this cancer, directly establishes super-enhancers of each of these three TFs to activate their transcription. In turn, KLF15, TCF4 and NKX2-2 co-bind to their own and each other’s super-enhancers and promoters, forming an inter-connected auto-regulatory loop. Functionally, CRC factors contribute significantly to cell proliferation of Ewing sarcoma both *in vitro* and *in vivo*, and are all overexpressed in this cancer. Mechanistically, CRC factors exhibit prominent capacity of co-regulating the epigenome in cooperation with EWS-FLI1, occupying 77.2% of promoters and 55.6% of enhancers genome-wide. Downstream, CRC TFs coordinately regulate gene expression networks in Ewing sarcoma, directly controlling important signaling pathways for cancer, such as lipid metabolism pathway, PI3K/AKT and MAPK signaling pathways. Together, molecular characterization of the oncogenic CRC model advances our understanding of the biology of Ewing sarcoma. Moreover, this study identifies CRC-downstream genes and signaling pathways, which may contain potential targets for therapeutic intervention for this malignancy.

## Introduction

As a developmental cancer carrying few genetic alterations, Ewing sarcoma is the second most common malignancy of bone and soft tissue predominantly occurring in adolescents and young adults (1). EWS-FLI1, the primary fusion driver, rewires fundamentally the transcriptome of Ewing sarcoma cells (2–5). Nevertheless, in many cases, EWS-FLI1 requires the cooperation of other transcriptional cofactors, such as WDR5 and CBP/p300, to regulate chromatin modification and gene expression (6–8). Moreover, EWS-FLI1-targeting transcription factors (TFs), such as MEIS1, NKX2-2, SOX2, and OTX2 (7, 9–11), are required for the fusion driver to fulfil its oncogenic function in Ewing sarcoma cells. Thus, despite having immense capacity in chromatin regulation, EWS-FLI1 still relies on additional cooperators and mediators to orchestrate gene expression programs in Ewing sarcoma. Hence, comprehensive and unbiased characterization of such partners and mediators is critical for further understanding the biology of Ewing sarcoma.

TFs coordinate cell type-specific transcriptional programs typically through regulating distal cis-regulatory elements, including enhancers and super-enhancers. Although hundreds of TFs are expressed to some extend in any given cell type (12), only a handful appear to be critical for establishing and maintaining cell type-specific gene expression network (13–15). Notably, this small set of TFs (sometimes called master TFs) often form a “Core Regulatory Circuitry (CRC)” (16–18), wherein each TF not only self-regulates but also co-regulates each other by directly co-binding to their super-enhancers. Although such CRC transcriptional paradigm has been identified in both normal and neoplastic cell types (19, 20), its functionality seems to be particularly critical for developmental tumors in children and young adults who carry few somatic genomic alterations. Indeed, some of the best characterized CRC models are established from neuroblastoma, medulloblastoma, as well as PAX3-FOXO1^+^ rhabdomyosarcoma, all of which have “quiet” genomic landscapes compared with adulthood tumors (21–23). For example, in MYCN-amplified neuroblastoma, a set of master TFs (HAND2, ISL1, PHOX2B, GATA3 and TBX2) assemble a functional CRC to orchestrate the unique gene expression program in this cancer type (22). Moreover, in PAX3-FOXO1-driven rhabdomyosarcoma, the fusion protein exploits super-enhancers to set up a CRC machinery in collaboration with the master TFs (MYOG, MYOD and MYCN); this CRC is addicted by rhabdomyosarcoma cells for survival and proliferation (23). Considering that Ewing sarcoma resembles these developmental cancers in terms of having both few genomic alterations and a single epigenomic driver, we postulated that such oncogenic CRC model might exist in Ewing sarcoma, to cooperate with the transcriptional function of EWS-FLI1. The current study was aimed to identify and characterize this CRC apparatus in Ewing sarcoma, and to elucidate its functional significance in this cancer.

## Results

### Identification of CRC under the regulation of EWS-FLI1 in Ewing sarcoma

We recently characterized the super-enhancer landscape in Ewing sarcoma and confirmed the essential role of EWS-FLI1 in regulating the epigenome of this cancer (11). As introduced earlier, considering that CRC is particularly important for childhood developmental cancers having few genomic lesions, we postulated that such CRC model might also contribute to regulating Ewing sarcoma transcriptome through either cooperating with or mediating the function of EWS-FLI1. To test this hypothesis, we first sought to identify mathematically master TFs with high inter-connectivity through binding to their super-enhancers by motif scanning using our established method (18, 24, 25). Because of the central role of EWS-FLI1 in establishing the enhancer landscape in Ewing sarcoma, we made significant modifications of the method by requiring that all candidate TFs have both EWS-FLI1 binding motif and binding peaks in their assigned super-enhancers. We initially focused on the A673 cell line, since it is a well-characterized Ewing sarcoma line with available H3K27ac and EWS-FLI1 ChIP-Seq results (7). As a result, a small set (n=9) of CRC candidates were identified (Fig. 1A), including NKX2-2 and FOS which are known functional cooperators of EWS-FLI1 (7, 26). Because CRC factors have high and specific expression in their corresponding cell types, we interrogated the Cancer Cell Line Encyclopedia (CCLE) dataset and noted that, compared with other 5 CRC candidates (NFATC2, FOS, IRF2, ZBTB7B and MEF2D, Supplement Fig. 1), the expression of 4 candidate TFs (KLF15, NKX2-2, TCF4 and RREB1) showed restricted expression pattern in Ewing sarcoma cell lines (defined as top 5 among all cell types, Fig. 1B). However, it should be noted that the specificity of RREB1 expression was overall poor, as most cancer types had comparable mRNA levels. As a parallel analysis, Pearson correlation coefficient of the mRNA levels of these 9 candidates was determined, considering the co-regulatory relationship between CRC members. Notably, the same set of 4 factors (KLF15, NKX2-2, TCF4 and RREB1) displayed strong positive correlations with each other (Fig. 1C). In contrast, this correlation pattern was much weaker in other non-Ewing sarcoma bone cancer cell lines (Fig. 1C, right panel).

**Figure 1.**
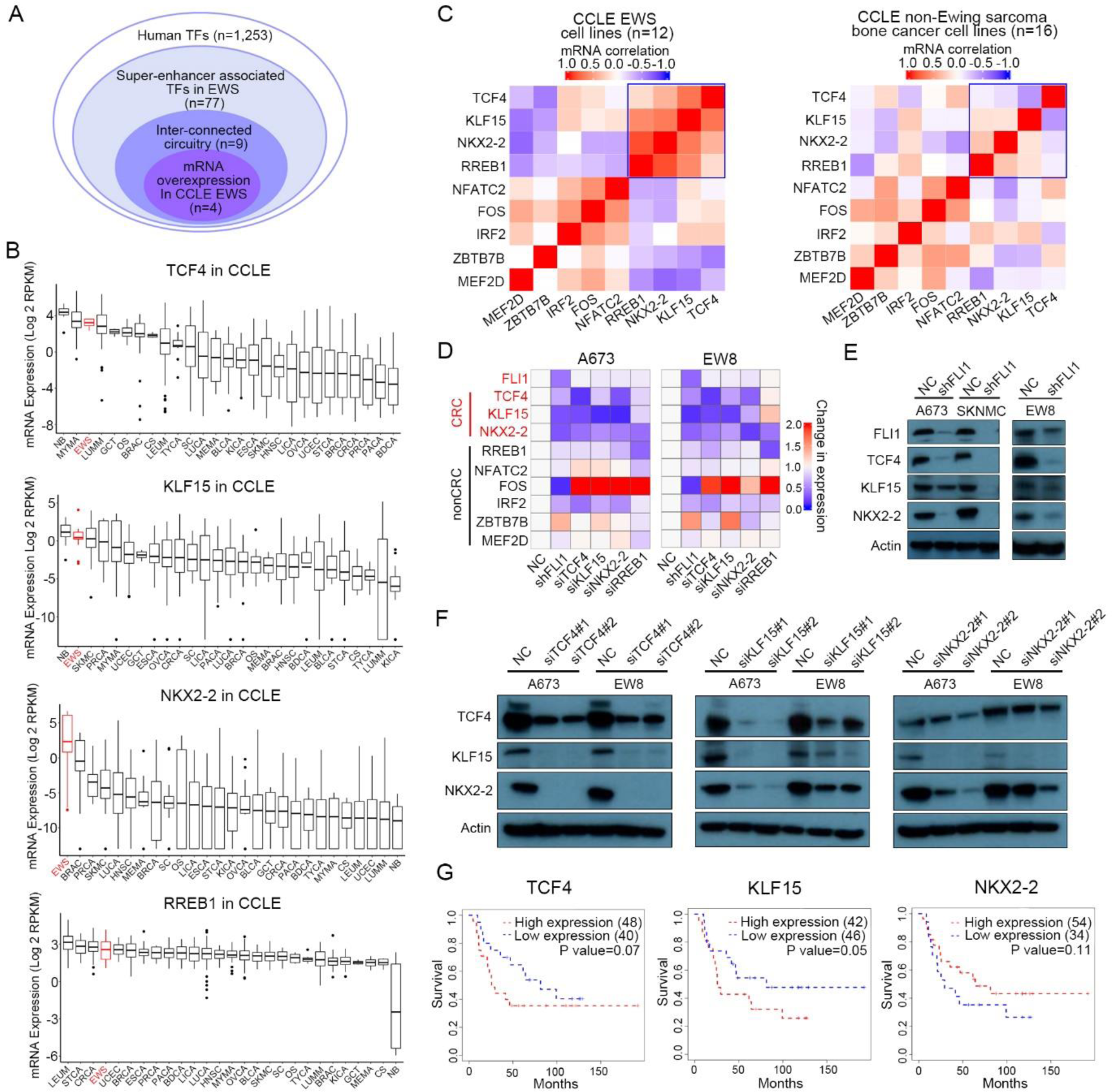
KLF15/TCF4/NKX2-2 form interconnected co-regulatory circuitry in Ewing sarcoma. (A) Integrative methods for identification of candidate CRC TFs. (B) The mRNA levels of candidate TFs in CCLE database. (C) Heatmap of Pearson correlation coefficient between candidate CRC TFs in Ewing sarcoma cell lines (n=12) or other bone cancer, but non-Ewing sarcoma, cell lines (n=16). Data were retrieved from CCLE database. (D) Heatmap of fold changes of mRNA expression of candidate TFs, following knockdown of either EWS-FLI1 or each of candidate TF in A673 and EW8 cells. (E) Immunoblotting of protein expression of CRC TFs upon either silencing of EWS-FLI1 or (F) silencing of each individual CRC TFs in A673 and EW8 cells. (G) Kaplan-Meier survival plots analyzing the expression of CRC TFs in Ewing sarcoma patients.

Based on these 4 candidates, we next sought to reconstruct functionally the CRC model and to determine their regulatory relationship with EWS-FLI1, by silencing of either EWS-FLI1 or each candidate individually. Knockdown of EWS-FLI1 strongly decreased mRNA expression of KLF15, NKX2-2 and TCF4, but not RREB1 (Fig. 1D). In contrast, the opposite effect was not observed, i.e., knockdown of candidate TFs had no impact on the level of EWS-FLI1, suggesting that the CRC is under the control of EWS-FLI1. Within the 4 CRC candidates, knockdown of either KLF15, NKX2-2 or TCF4 decreased the expression of the other two (Fig. 1D), but not RREB1. On the other hand, RREB1-silencing did not produce these co-regulatory effects, demonstrating that in Ewing sarcoma cells, KLF15, TCF4 and NKX2-2 together constitute an inter-connected circuitry. Importantly, all of these regulatory effects were validated at the protein levels with either additional shRNA or siRNAs in both A673 and EW8 cell lines (Figs. 1E, 1F). In a univariate analysis, high expression of either KLF15 or TCF4 in Ewing sarcoma was associated with significantly poor survival (Fig. 1G).

### EWS-FLI1 directly activates three CRC factors

To elucidate the co-regulatory mechanisms between CRC TFs and how they are controlled by EWS-FLI1, ChIP-Seq was performed using specific antibodies against either KLF15, TCF4 or NKX2-2 in A673 cells. Importantly, these 3 TFs trio-occupied both super-enhancers and promoters of each other, as well as themselves (Fig. 2A, Supplement Fig. 2, 3), forming an interconnected circuitry as predicted by our method. Enrichment of H3K27ac signals at these super-enhancers was observed across all Ewing sarcoma primary tumor samples (n=3) and cell lines (n=4). All three distal super-enhancers, but not promoters, were occupied by EWS-FLI1 in both A673 and SKNMC cells (Fig. 2A, Supplement Fig. 2), again validating our method. Furthermore, by re-analyzing the publicly-available Hi-C data of SKNMC cells, we confirmed interactions between super-enhancers with promoters in every gene locus, suggesting direct transcriptional co-regulation between KLF15, TCF4 and NKX2-2 by binding to each other’s super-enhancers. These data also highlight that EWS-FLI1 directly controls the expression of all three CRC members by occupying their super-enhancers (Fig. 2A).

**Figure 2.**
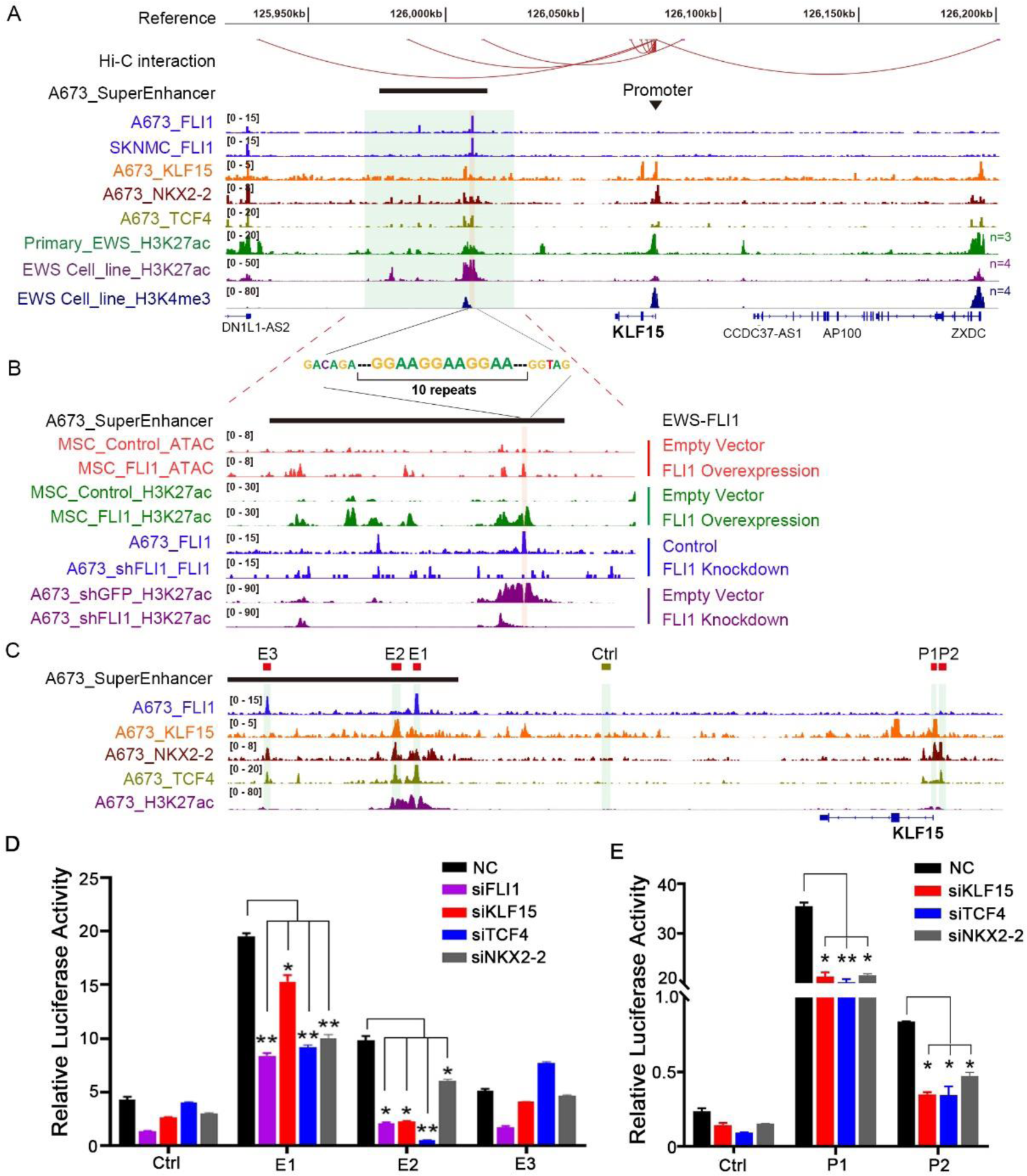
EWS-FLI1 and CRC TFs co-operatively activate the transcription of KLF15. (A) Integrative Genomics Viewer (IGV) plots of ChIP-Seq showing co-occupancy of EWS-FLI1 and CRC TFs at the super-enhancer and promoter of the KLF15 gene locus. Hi-C interactions were re-analyzed from the Hi-C data of SKNMC cell line downloaded from ENCODE database; H3K27ac, H3K4me3 and EWS-FLI1 ChIP-Seq data were retrieved from GEO (GSE61953). (B) ATAC-Seq and ChIP-Seq profiles at KLF15 super-enhancer region in either the presence or absence of EWS-FLI1 overexpression (top 4 tracks) or knockdown (bottom 4 tracks). Data were retrieved from GEO (GSE61953). (C) Zoom in view of ChIP-Seq signals in KLF15 locus. Three putative enhancer elements (E1-E3), two promoter elements (P1, P2) and one negative control region were separately cloned into luciferase reporter vectors. (D) Enhancer and (E) promoter activities were measured by luciferase reporter assays in A673 cells in either the presence or absence of knockdown of indicated TFs. Mean ± s.d. are shown, n = 3. *, P < 0.05; **, P < 0.01.

Previous *de novo* motif analyses elucidate that cis-regulatory elements activated by EWS-FLI1 are strongly enriched for GGAA repeats (7). Consistently, prominent enrichment of GGAA repeats were observed in EWS-FLI1 binding sites at all three CRC TFs (Figs. 2A-B, Supplement Fig. 2). To further examine whether and how EWS-FLI1 establishes the super-enhancers of CRC factors, we first interrogated chromatin accessibility of normal primary MSCs infected with an EWS-FLI1 expression vector. In the absence of ectopic EWS-FLI1 expression, these genomic regions had negligible ATAC-Seq peaks or H3K27ac signals, indicating little transcriptional activity. Importantly, ectopic expression of EWS-FLI1 converted these regions to super-enhancers with much higher H3K27ac intensity and chromatin accessibility (top 4 tracks in Fig. 2B, Supplement Fig. 2). On the other hand, in A673 cells, knockdown of EWS-FLI1 drastically decreased H3K27ac signal in EWS-FLI1-occupied super-enhancers (bottom 4 tracks in Fig. 2B, Supplement Fig. 2). These results strongly suggest that EWS-FLI1 directly initiates and maintains the super-enhancers of CRC members.

Next, we identified three enhancer constituents (E1, E2, E3) within the super-enhancer of KLF15 based on the occupancy of TFs (Fig. 2C). Specifically, E1 and E3 were trio-occupied by EWS-FLI1, TCF4 and NKX2-2, while E2 was trio-occupied by KLF15, TCF4 and NKX2-2 (Figs. 2A, 2C). These enhancer elements, as well as a control region, were subsequently cloned into the pGL3-promoter luciferase reporter vector which were transfected into A673 cells. Robust reporter activities of E1 and E2, but not E3, were observed (Fig. 2D). To determine the regulation of these enhancers by their occupying TFs, we silenced each TFs, and knockdown of each factor (Fig. 2D) markedly reduced the activity of E1 and E2. These results support direct regulation of KLF15 super-enhancer by all CRC members, as well as EWS-FLI1. Moreover, CRC factors, but not the fusion oncoprotein, co-bound the promoter region of KLF15 (Fig. 2A). We similarly cloned two promoter constituents of KLF15 (P1 and P2) into the pGL3-basic luciferase reporter vector, and measured their activities by reporter assays in A673 cells (Fig. 2E). Both of the two promoter regions expectedly showed strong reporter activity, and they were significantly inhibited upon silencing each of the three CRC TFs.

### CRC factors cooperate with EWS-FLI1 to orchestrate the transcriptional network of Ewing sarcoma cells

To gain insights into the mechanistic basis of CRC TFs in regulation of the transcriptome of Ewing sarcoma, we analyzed the epigenomic characteristics of their occupancy in A673 cells. To this end, we first annotated the putative promoter (H3K4me3^+^/H3K27ac^+^/H3K4me1^-^) and distal enhancer (H3K4me3^-^/H3K27ac^+^/H3K4me1^+^) regions by available histone marks. Genome-wide peaks of each CRC TFs and EWS-FLI1 were then assigned to these regions. Notably, the occupancy of these TFs was pervasive throughout the genome, with 77.2% of all promoters and 55.6% of all enhancers bound by at least one of the factors (Fig. 3A). Although genomic regions quadruple-bound by all 4 TFs were rare (0.6% promoters and 0.4% enhancers), cooperative occupancy (i.e., trio- and dual-occupation) were more common than solo-occupancy in both the promoter and enhancer regions (Figs. 3A, 3B, Supplement Fig. 3), suggesting strong cooperativity between these factors. Specifically, several important co-binding patterns were observed: i) Validating previous studies (6, 7), EWS-FLI1 peaks were restricted within enhancer regions; ii) EWS-FLI1 almost always trio-occupied with TCF4/NKX2-2; iii) KLF15-binding was restricted to promoters, and it either dual-occupied with TCF4 or trio-occupied with TCF4/NKX2-2.

**Figure 3.**
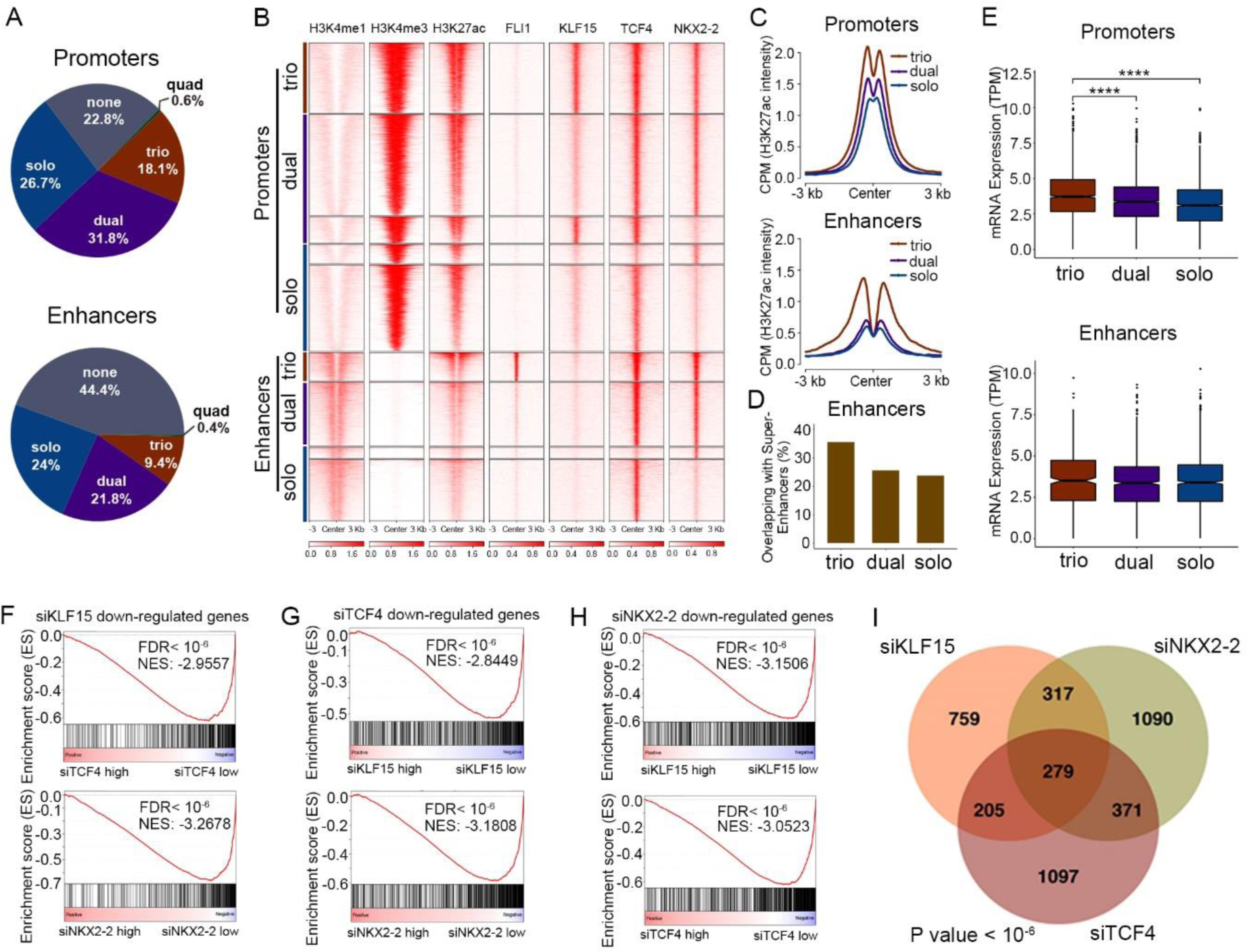
Transcriptional cooperativity between EWS-FLI1 and CRC TFs. (A) Pie charts of the factions of combinatorial binding patterns of EWS-FLI1 and three CRC TFs. (B) Heatmaps of ChIP-Seq signals of indicated factors in A673 cells, stratified by different combinatorial binding patterns. (C) Line plots of H3K27ac ChIP-Seq signals from indicated groups of peaks in A673 cells. (D) The overlapping of indicated groups of peaks with super-enhancers in A673 cells. (E) Box plots of mRNA expression of genes associated with indicated groups of peaks in A673 cells. (F) GSEA plots showing the enrichment of downregulated genes upon knockdown of KLF15 in either siTCF4 or siNKX2-2 RNA-Seq data. Reciprocal GSEA plots of RNA-Seq following knockdown of TCF4 (G) and NKX2-2 (H) are similarly provided. NES, normalized enrichment score; FDR, false discovery rate. (I) Venn diagram of downregulated genes following silencing of each single TFs (p value < 10^-6^; empirical distribution test).

Importantly, regardless of promoter or enhancer elements, trio-binding regions always harbored much higher H3K27ac intensity compared to either solo- or dual-binding regions (Fig. 3C), suggesting that regulatory regions trio-occupied by these TFs may have stronger transcriptional activity. Moreover, tri-occupied regions were more likely to overlap super-enhancers than either dual- or solo-occupied regions (Fig. 3D).

To determine further the possible transcriptional impact of these binding event, matched RNA sequencing (RNA-Seq) data of A673 cells were analyzed. Notably, genes associated with trio-bound promoters exhibited the highest expression levels compared with either solo-or dual-bound promoters (Fig. 3E, upper panel). A similar trend was also observed in enhancer elements, albeit without statistical significance (Fig. 3E, lower panel). Together, these data highlight the cooperative binding as a key characteristic of CRC TFs. Moreover, CRC factors cooperate not only among themselves, but also with EWS-FLI1 in regulating the epigenome of Ewing sarcoma cells.

To establish further the regulatory effect of CRC factors on the transcriptome, we performed RNA-Seq of A673 cells in either the presence or absence of knockdown of each TF. Importantly, gene set enrichment analyses (GSEA) showed that genes decreased following silencing of KLF15 were strongly and significantly enriched in those also downregulated upon depletion of either TCF4 or NKX2-2 (Fig. 3F). The same pattern was also observed in RNA-Seq data of either siTCF4 or siNKX2-2 (Figs. 3G, 3H). Indeed, the downregulated genes in each dataset substantially and significantly overlapped (p < 10^-6^, empirical distribution test); in fact, the “shared” changes accounted for almost half of all changes in every case (43.8% for TCF4, 51.3% for KLF15, and 47.0% for NKX2-2) (Fig. 3I). These results strongly suggest that CRC factors act in concert with each other to co-regulate the transcriptome of Ewing sarcoma cells.

### CRC factors have strong tumor-promoting function in Ewing sarcoma cells

Considering the prominent roles of CRC in controlling transcriptional network (27, 28), we postulated that KLF15, TCF4 and NKX2-2 are required for the viability and proliferation of Ewing sarcoma cells. Indeed, NKX2-2 has been established as a tumor-promoting factor in Ewing sarcoma (29), but the functions of KLF15 and TCF4 remained hitherto unknown in this cancer. We thus performed loss-of-function assays and showed that knockdown of either KLF15, TCF4 or NKX2-2 by individual siRNAs (Fig. 4A) markedly inhibited cell proliferation (Fig. 4B, Supplement Fig. 4A,) and colony growth (Fig. 4C). The results were verified by doxycycline-inducible expression of multiple independent shRNAs (Supplement Figs. 4B-4E). Fluorescence-activated cell sorting (FACS) analysis showed that depletion of endogenous expression of CRC TFs caused cell cycle arrest (Supplement Figs. 4F, 4G). We also detected increased cell apoptosis after knockdown of TCF4 (Supplement Fig. 4H). Additionally, silencing of KLF15 decreased cell migration of Ewing sarcoma cells and its over-expression produced the opposite effect (Supplement Figs. 4I-4O). A previous study showed that NKX2-2 silencing suppressed xenograft growth of Ewing sarcoma (30, 31). Here, we tested the dependency of Ewing sarcoma cells on KLF15 and TCF4 *in vivo*. Expression of doxycycline (DOX)-inducible shRNAs against either KLF15 or TCF4 potently inhibited xenograft growth of Ewing sarcoma (Figs. 4D, 4E). Finally, immunoblotting of xenograft tumors again confirmed the co-regulation of CRC TFs, as shown by the downregulation of each CRC member upon silencing of either KLF15 or TCF4 (Fig. 4F). Taken together, these results demonstrate that as CRC members, KLF15, TCF4 and NKX2-2 positively co-regulate each other and promote the growth and survival of Ewing sarcoma cells.

**Figure 4.**
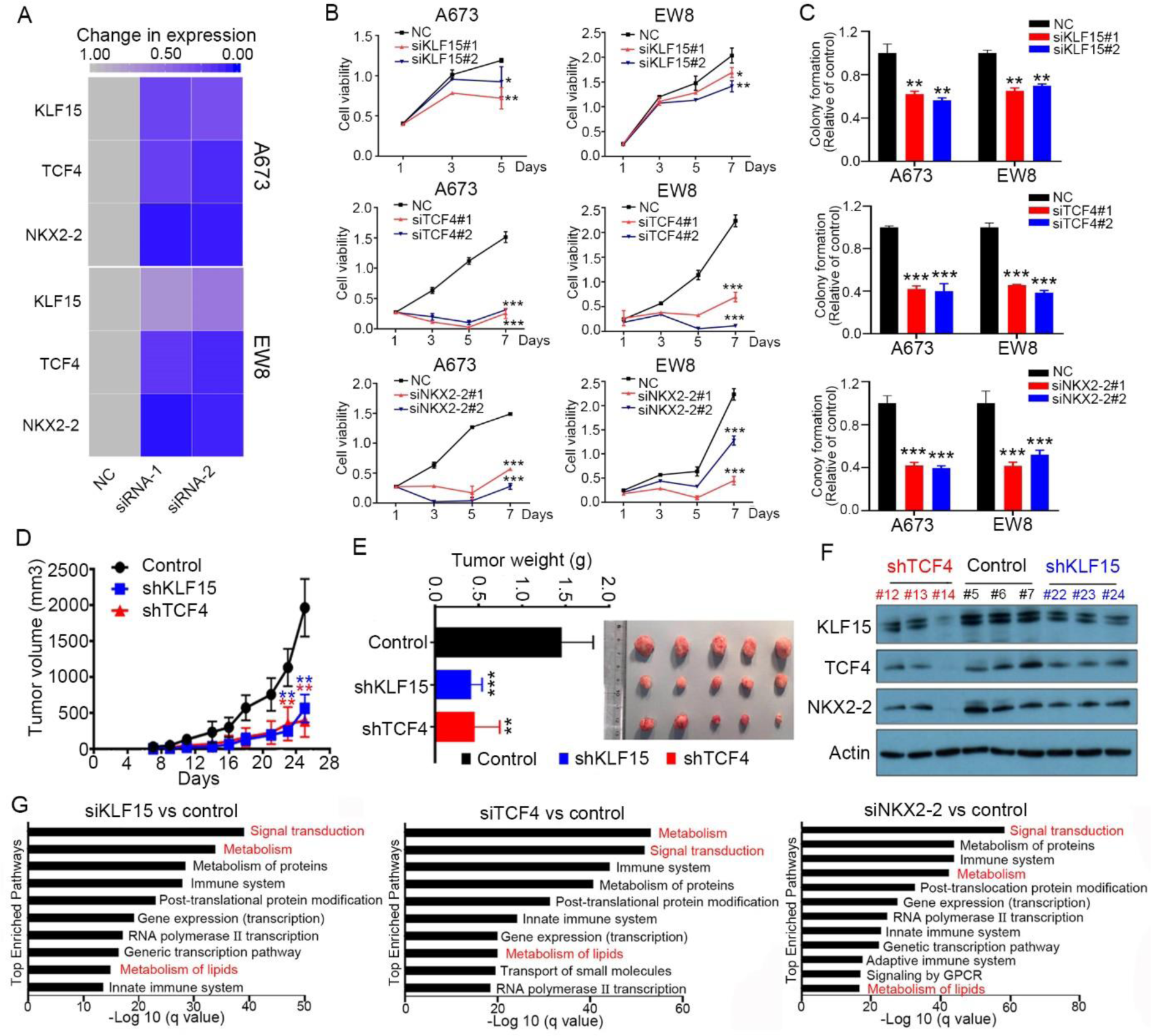
Cancer-promoting functions of CRC TFs in Ewing sarcoma cells. (A) Heatmap of mRNA levels of three TFs before and after knockdown by individual siRNAs in A673 and EW8 cells. (B-C) Silencing of three TFs by individual siRNAs decreased cell proliferation (B) and colony growth (C) in Ewing sarcoma cells. (D-F) Silencing of either KLF15 or TCF4 by inducible shRNAs decreased xenograft growth *in vivo*. Growth curves (D), tumor weights and images (E), and co-regulation between CRC TFs (F) in resected tumors. Mean ± s.d. are shown, n = 6. **, P < 0.01; ***, P < 0.001. (G) Pathway enrichment analysis of downregulated genes after knockdown of each of three CRC TFs in A673 cells.

### CRC factors co-regulate lipid metabolism pathways in Ewing sarcoma

Having established the functional significance of CRC TFs in the proliferation and viability of Ewing sarcoma cells, we next focused on investigating their downstream pathways. Pathway enrichment analysis was performed using downregulated genes, defined as log 2 (fold change) < -0.5 and q value < 0.05, upon knockdown of each TF individually. Multiple top enriched terms were shared among each analysis, including “Signal transduction”, “Metabolism”, “Metabolism of lipids”, “Gene expression (transcription)” and “RNA Pol-II transcription” (Fig. 4G), strongly supporting the functional cooperation between these CRC factors. Among these shared top-ranking pathways, we were particularly interested in the lipid metabolism signaling, since: i) the biological significance of lipid metabolism in the biology of Ewing sarcoma is unknown; ii) the regulatory basis of CRC TFs on lipid metabolism is unclear.

To elucidate the mechanisms underlying the regulation of lipid metabolism by CRC TFs, we analyzed in-depth all the enriched genes (n=242) in the term “Metabolism of lipids” (Fig. 5A). Again, consistent with the close cooperation between CRC TFs, these genes were strongly shared between individual RNA-Seq data upon knockdown of each CRC factors. These downregulated genes were enriched in key pathways in lipid metabolism, including glycerophospholipid metabolism, sphingolipid metabolism, steroid biosynthesis, biosynthesis of unsaturated fatty-acids, as well as fatty-acid elongation (Fig. 5B). To quantify the functional impact of CRC TFs on these lipid metabolism processes, we performed liquid chromatography tandem mass spectrometry (LC-MS/MS)-based lipidomics in A673 cells, considering the immense structural complexity of lipid species. Specifically, quantitative lipidomics were performed in the presence and absence of the knockdown of each individual TF. To warrant reproducibility and robustness, 3-4 replicates of each sample were measured, and the data were highly comparably and consistent (Supplement Fig. 5). In total, we identified an average of 1,591 lipid ions in A673 cells, which belonged to 30 lipid classes, suggesting high sensitivity and lipidome coverage of the methodology. The most abundant lipid classes in A673 cells were phosphatidylcholine (PC), phosphatidylethanolamine (PE), diacylglycerol (DG), triacylglycerol (TG), dimethylphosphatidylethanolamine (dMePE) and sphingomyelin (SM). The most abundant fatty acyl chains were oleate (C18:1), palmitate (C16:0), stearate (C18:0), palmitoleic (C16:1), arachidonate (C20:4), docosahexaenoic (C22:6). In agreement with the pathway enrichment analysis, silencing of each TF caused appreciable changes in the lipid landscape of A673 cells (Figs. 5C-5E). Specifically, comparing control with knockdown groups, 251 (in siKLF15), 397 (in siTCF4) and 319 (in siNKX2-2) lipid ions were differentially regulated (q value < 0.05, absolute fold change > 2). Although the alterations in lipid species were variable between different knockdown experiments, the majority of which were notably converged to two lipid-associated pathways: glycerophospholipid and sphingolipid pathways (Fig. 5F). These changes in lipid classes were highly consistent with and supportive of our pathway analyses of RNA-Seq data, which identified the top two enriched pathways as glycerophospholipid metabolism and sphingolipid metabolism (Fig. 5B).

**Figure 5.**
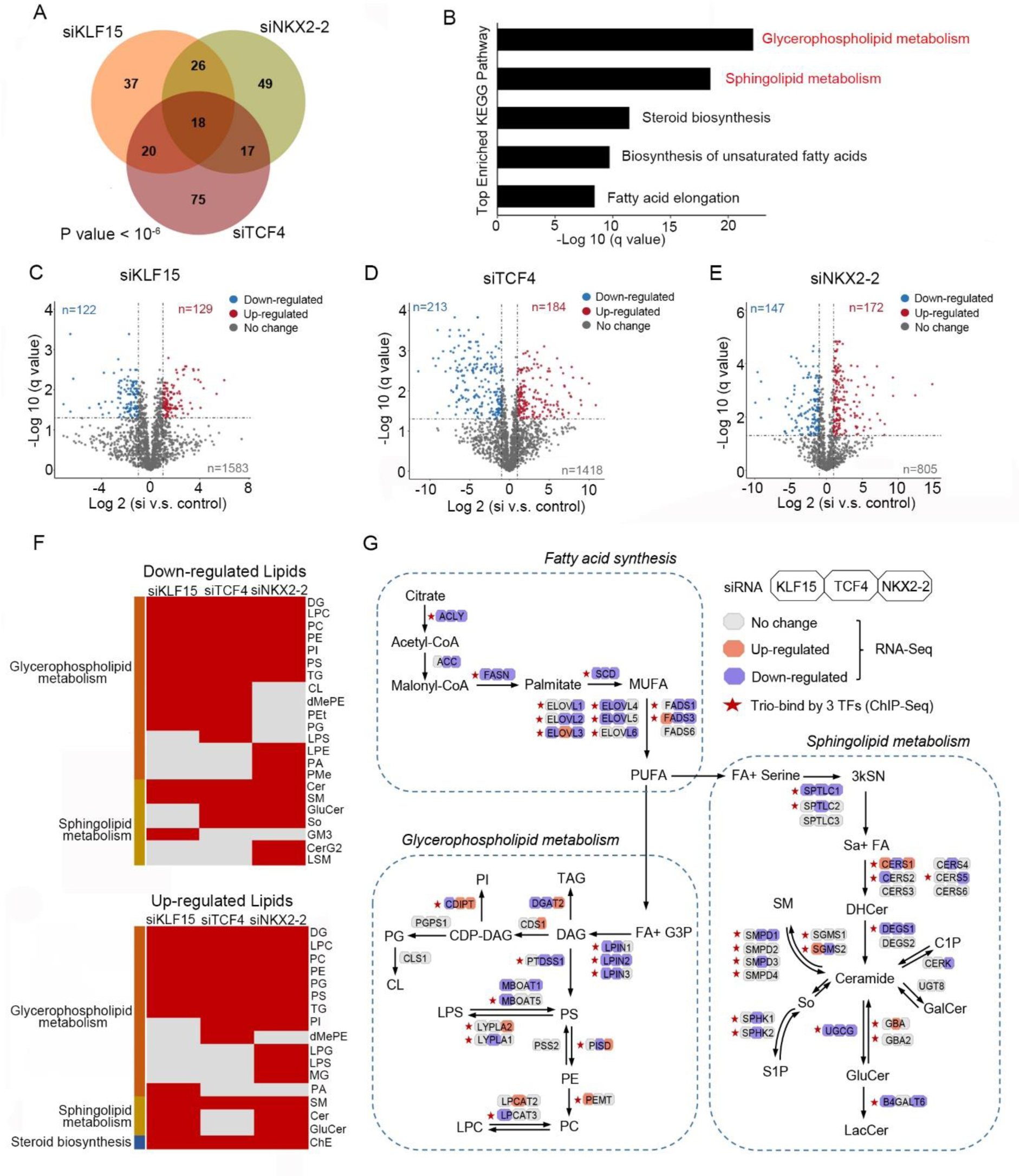
CRC TFs regulate lipid metabolism in Ewing sarcoma. (A) Venn diagram and (B) KEGG pathway enrichment of downregulated lipid metabolism-associated genes following silencing of each single CRC TFs (p value < 10^-6^; empirical distribution test). (C-E) Volcano plots of lipidomic analyses showing differentially regulated lipid ions upon silencing of either (C) KLF15, (D) TCF4 or (E) NKX2-2. Each dot is a lipid ion. (F) Heatmaps of alterations in lipid classes upon silencing of each CRC TFs. Glycerophospholipid metabolism associated lipid classes: DG, LPC, PC, PE, TG, PG, PS, PI, PA, LPG, LPS, MG, dMePE, LPE, PMe, CL and PEt; Sphingolipid metabolism associated lipid classes: SM, GluCer, Cer, So, GM3, CerG2 and LSM; Steroid biosynthesis associated lipid classe: ChE. (G) A diagram showing lipid metabolism which was perturbed by silencing of CRC TFs via integration of RNA-Seq, ChIP-Seq and lipidomic results. Color-coded bars indicate relative expression changes in RNA-Seq after silencing of each of the three TFs.

We next examined in detail how lipid metabolism was perturbed by silencing of CRC TFs via integration of RNA-Seq, ChIP-Seq and lipidomic results. This systematic approach identified many important lipid enzymes as direct targets of CRC TFs. For example, rate-limiting enzymes for fatty-acid synthesis and elongation (FASN, ACLY and SCD) were trio-occupied and directly regulated by KLF15, TCF4 and NKX2-2. Moreover, sphingolipid anabolic enzymes (including SPTLC1, DEGS1 and UGCG) and glycerophospholipid anabolic enzyme (LPIN2) were under direct co-regulation by all three CRC TFs (Fig. 5G). Not surprisingly, we also observed solo-or dual-regulation by only 1 or 2 TFs on many other enzymes, such as CERS2 (by KLF15), CERS4 (by TCF4), CERS5 (by NKX2-2) and B4GALT6 (by both KLF15 and NKX2-2). These findings together suggest that CRC factors co-operatively and directly regulate lipid metabolism pathways in Ewing sarcoma cells.

### Biological significance of lipid metabolism in Ewing sarcoma cells

Following the characterization of the profound co-regulatory effects of CRC factors on lipid metabolic processes, we next sought to explore the biological significance of lipid metabolism in Ewing sarcoma cells. We first tested rate-limiting enzymes for fatty-acid synthesis (FASN, ACLY and SCD, which were trio-regulated by three CRC TFs), since this is the initial process upstream of all lipid metabolism reactions. Verifying our RNA-Seq results, qRT-PCR showed knockdown of each single TF reduced the expression of these central enzymes (Fig. 6A). Immunoblotting confirmed downregulation at the protein levels of FASN, ACLY and SCD (Fig. 6B). In addition, silencing of TCF4 (but not KLF15 or NKX2-2) also reduced the expression of ACC, another key enzyme in fatty-acid synthesis. Some of the trio-binding peaks were shown in Fig. 6C using *FASN* and *SCD* promotors as examples.

**Figure 6.**
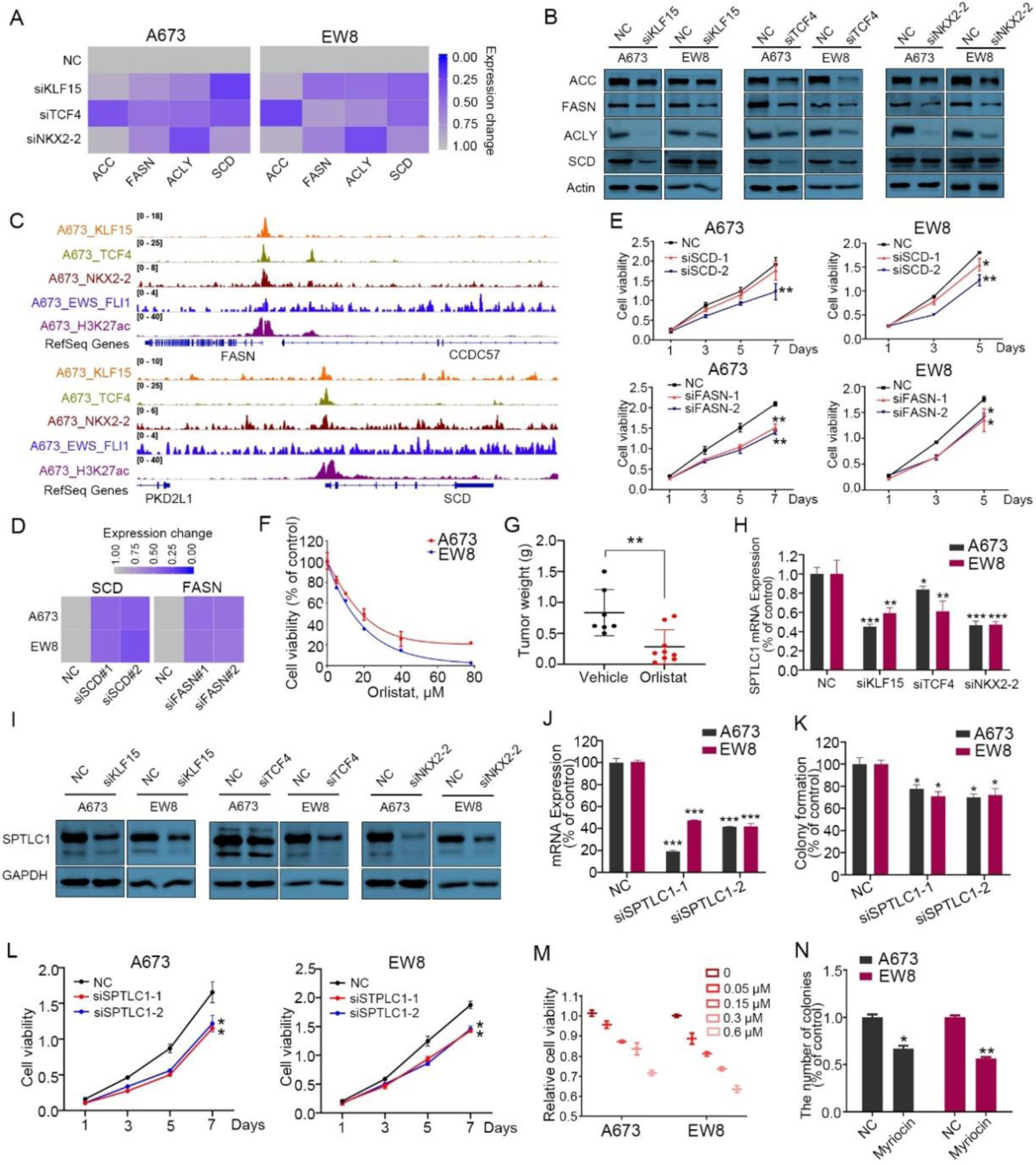
Biological functions of lipid synthesis pathway in Ewing sarcoma cells. (A, B) Silencing of each of three CRC TFs decreased the expression of lipid biosynthetic enzymes at both the mRNA (A) and protein levels (B) in Ewing sarcoma cell lines. (C) IGV plots showing *FASN* and *SCD* promoters which were trio-occupied by three CRC TFs. (D) Silencing of FASN and SCD by siRNAs (E) decreased Ewing sarcoma cell proliferation. (F) *In vitro* MTT proliferation assay of Ewing sarcoma cells in the presence of different doses of orlistat (FASN inhibitor). (G) Orlistat treatment suppressed xenograft growth *in vivo*. Weights of resected tumors from both groups are shown. (H, I) Silencing of each CRC TFs decreased SPTLC1 expression at both the mRNA (H) and protein levels (I) in Ewing sarcoma cell lines. (J-L) Silencing of SPTLC1 by siRNA (J) decreased colony growth (K) and cell proliferation (L) in Ewing sarcoma cell lines. (M, N) SPTLC inhibitor (Myriocin) decreased cell proliferation (M) and colony growth (N) in both A673 and EW8 cells. Data are presented as mean±SD of three replicates. *, P < 0.05; **, P < 0.01; ***, P < 0.001.

Importantly, silencing of either FASN or SCD markedly reduced cell proliferation and colony growth of both A673 and EW8 cell lines (Figs. 6D, 6E). Moreover, Orlistat, a specific FASN inhibitor, potently inhibited cell proliferation both *in vitro* (Fig. 6F) and *in vivo* (Fig. 6G), suggesting the biological importance of fatty-acid synthesis pathway in Ewing sarcoma cells.

In addition to fatty-acid synthesis, we also explored the functional significance of sphingolipid metabolism, another top-enriched lipid pathway in both RNA-Seq and lipidomics (Figs. 5B, 5G). Among all the sphingolipid anabolic enzymes regulated by CRC, SPTLC1 acts as the rate-limiting enzyme for *de novo* sphingolipid biosynthesis (32). Indeed, knockdown of each single TF inhibited the expression of SPTLC1 at both transcriptional and protein levels (Figs. 6H, 6I). Importantly, silencing of SPTLC1 reduced both colony growth and cell proliferation of Ewing sarcoma cells (Figs. 6J-6L). Moreover, treatment of Myriocin, a SPTLC1 inhibitor, reduced cell proliferation (Fig. 6M) and colony growth (Fig. 6N) of both A673 and EW8 cells.

### CRC TFs co-regulate PI3K/AKT and MAPK signaling pathways

In parallel, we also investigated in-depth the term “Signal transduction” since it was ranked highest in all three RNA-Seq upon knockdown of CRC TFs (1^st^ in both siKLF15 and siNKX2-2 and 2^nd^ in siTCF4, Fig. 4G). To dissect which specific signaling pathways were coordinately regulated by CRC TFs, KEGG pathway enrichment was next performed using the genes enriched in “Signal transduction” term (the union set, n=669) following silencing of each factor (Fig. 7A). Notably, PI3K/AKT and MAPK signaling pathways were identified as the top-ranked pathways (Fig. 7A). Focusing on these two pathways, we found that genes downregulated upon knockdown of each CRC factors were strongly overlapped. Specifically, 32/84 (38%) genes in PI3K/AKT signaling and 23/72 (32%) genes in MAPK signaling were decreased in at least two out of three knockdown groups (Fig. 7B), supportive of the notion that CRC factors cooperatively regulate these pathways.

**Figure 7.**
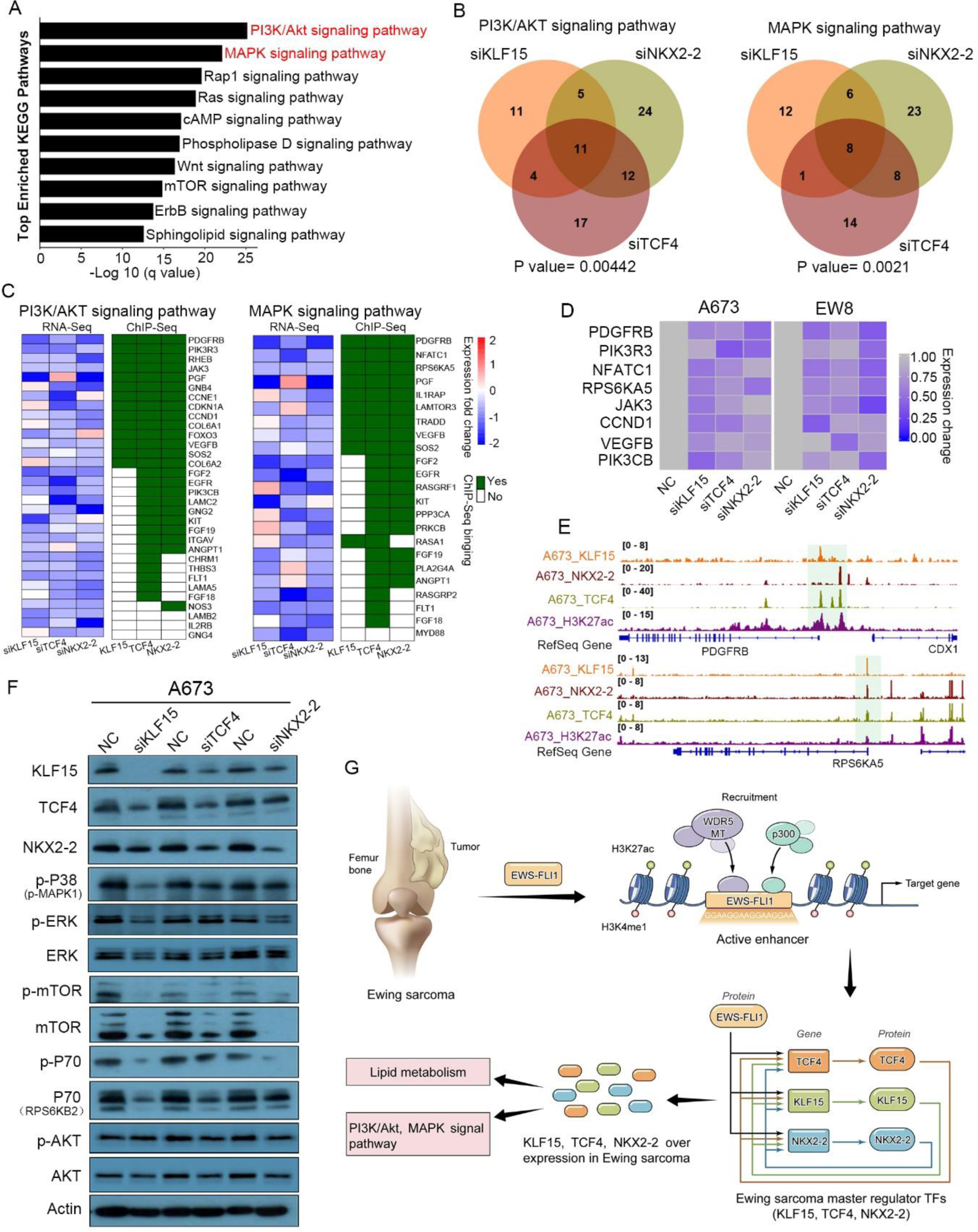
PI3K/AKT and MAPK signal pathways are regulated by CRC TFs in Ewing sarcoma cells. (A) KEGG pathway enrichment of downregulated genes after silencing each of the three CRC TFs. (B) Venn diagrams of downregulated genes in PI3K/AKT (left panel) and MAPK (right panel) pathways following silencing of each single CRC TFs. (C) Heatmaps showing the mRNA expression and binding patterns of the downregulated genes of these two pathways in at least two knockdown groups. (D) Silencing of each single CRC TFs inhibited the mRNA expression of key molecules of PI3K/AKT and MAPK signal pathways. (E) IGV plots showing *RPS6KA5* and *PDGFRB* promoters which were trio-occupied by three CRC TFs. (F) Immunoblotting assay showing the total and phosphorylation levels of key mediators of PI3K/AKT and MAPK pathways upon silencing of each of the three CRC TFs. (G) A proposed model of transcriptional dysregulation mediated by EWS-FLI1 and CRC TFs in the biology of Ewing sarcoma.

Further integration with KLF15, TCF4 and NKX2-2 ChIP-Seq data identified that 40/44 (90.9%) commonly downregulated genes (in at least two knockdown groups) of these two cascades were directly occupied by at least one of the CRC TFs (Fig. 7C). Moreover, 23 and 19 of these 40 target genes (57.5% and 47.5%) were dual- and trio-occupied by CRC TFs, respectively (Fig. 7C). This result suggests strong cooperation between CRC TFs in the regulation of these two signaling pathways. These direct co-targets included key mediators or effectors of PI3K/AKT and MAPK pathways, such as PDGFRB, PIK3R3, JAK3, VEGFB, RPS6KA5, CCND1 and NFATC1 (Fig. 7C). Indeed, silencing of either KLF15, TCF4 or NKX2-2 inhibited the expression of these key molecules as validated by qRT-PCR (Fig. 7D). Some of the trio-binding peaks were shown in Fig. 7E using the *RPS6KA5* and *PDGFRB* promoters as examples. Moreover, immunoblotting assays confirmed reduced phosphorylation levels of several central mediators of PI3K/AKT and MAPK pathways, including P38, ERK, mTOR, P70 and AKT (Fig. 7F). Taken together, these data demonstrate that KLF15, TCF4 and NKX2-2 coordinately promote the transcription of chief components of the PI3K/AKT and MAPK signaling pathways, thereby activating these pro-growth and pro-survival signaling cascades in Ewing sarcoma.

## Discussion

EWS-FLI1 is a major determinant of genome-wide chromatin states in Ewing sarcoma (2-5, 33, 34), by functioning as a pioneer factor (7, 9, 10, 35). Nevertheless, the fusion driver requires other TFs and cofactors to fulfill its oncogenic activities. Considering the essential role of CRC in other developmental cancers driven by single transcriptional regulators (such as NMYC-driven neuroblastoma and PAX3-FOXO1-driven rhabdomyosarcoma) (22, 23), we postulated that in Ewing sarcoma, a functional CRC apparatus might cooperate with EWS-FLI1 in the regulation of Ewing sarcoma transcriptome. In order to address this hypothesis, we modified a computational algorithm given the central role of EWS-FLI1, and identified a CRC “trio” constituted by KLF15, TCF4 and NKX2-2 in Ewing sarcoma cells.

Integrative epigenomics analyses of ChIP-Seq, ATAC-Seq as well as Hi-C together demonstrate that EWS-FLI1, as a pioneer factor, directly establishes the super-enhancers of each of the three CRC TFs to activate their transcription. Subsequently, KLF15, TCF4 and NKX2-2 co-bind to their own and each other’s super-enhancers (together with EWS-FLI1 binding) and promoters (without EWS-FLI1 binding), forming an inter-connected auto-regulatory loop (Fig. 7G).

In addition to cooperating with EWS-FLI1 to co-amplify their own transcription through a CRC model, KLF15, TCF4 and NKX2-2 also exhibit prominent capability of co-regulating the epigenome of Ewing sarcoma cells. In particular, the cooperative occupancy of CRC TFs is pervasive, covering the majority (77.2%) of all putative promoters and over half (55.6%) of all putative enhancers (Fig. 3A). Notably, TCF4 and NKX2-2 have higher capacity in occupying the genome than either EWS-FLI1 or KLF15, as their binding regions almost always encompass those of EWS-FLI1 and KLF15. Specifically, TCF4 and NKX2-2 serve as co-binding partners for EWS-FLI1 (in enhancer regions) and for KLF15 (in promoter regions), in addition to their own binding regions. Nevertheless, these three TFs still tend to operate collaboratively, such that their co-bindings events (trio- and dual-) are more common than solo-binding events. Moreover, cooperative binding by more TFs appears to be associated with higher transcriptional activity. Indeed, DNA regulatory elements loaded with more TFs always have higher H3K27ac intensity (Fig. 3C), which are also associated with higher expression of downstream genes (only significantly in promoter regions, Fig. 3E). These findings are consistent with the proposed function of combinatorial binding of multiple factors in close proximity, which is to overcome with more potency the energetic barrier for nucleosome eviction, thus facilitating activation of cis-regulatory elements.

Not surprisingly, the cooperative binding of CRC TFs results in co-regulation of gene expression program in Ewing sarcoma cells. Indeed, GSEA of RNA-Seq data confirms that downregulated genes upon knockdown of each TF strongly and significantly overlapped with each other. In fact, the shared changes account for almost half of all changes in every RNA-Seq of CRC TFs, strongly suggesting that KLF15, TCF4 and NKX2-2 coordinate to regulate the transcriptome of Ewing sarcoma cells (Figs. 3F-3I). Phenotypically, similar to CRC factors in other cancer types, KLF15, TCF4 and NKX2-2 each shows strong pro-growth and pro-survival functions in Ewing sarcoma. Indeed, NKX2-2 has been reported to promoting cell proliferation of Ewing sarcoma (29–31), however, the biological significance of either KLF15 or TCF4 had hitherto been unknown in this cancer.

TCF4 (also known as ITF2) is a basic helix-loop–helix (bHLH) TF that plays a crucial role in the differentiation and specification of central nervous system (CNS) (31, 36). Interestingly, depending on different tumor types, TCF4 functions as either an oncogene (diffuse large B-cell lymphoma) (37) or tumor suppressor (medulloblastoma and colon cancer) (38, 39). KLF15 is a key regulator of metabolic pathways controlling adipogenesis and gluconeogenesis in both liver and skeletal muscles (40, 41). However, its involvement in human malignancies has been poorly understood with mixed findings. For example, KLF15 has shown anti-proliferative activities in gastric and breast cancer cells (42, 43); however, KLF15 promotes the proliferation and metastasis of lung adenocarcinoma cells (44). These reports suggest that the cancer-specific functions of KLF15 and TCF4 are highly context-dependent. In our study, consistent with their Ewing sarcoma-promoting functions, all CRC factors are expressed particularly robustly in Ewing sarcoma relative to most of other cancer types (Fig. 1B).

To understand the functional contribution of CRC TFs in Ewing sarcoma, we performed pathway enrichment analyses of RNA-Seq upon knockdown of each factors. This approach further substantiates the cooperativity of KLF15, TCF4 and NKX2-2, since multiple top-enriched pathways are shared between three independent loss-of-function experiments. Two of the overlapped high-ranking pathways (lipid metabolism and signal transduction) were prioritized for in-depth investigation given their importance in cancer biology.

Dysregulated lipid metabolism is one of the most important metabolic hallmarks of cancer cells, which is pivotal for synthesis of cellular building blocks and signaling molecules (45–48). Accumulating evidence suggest that cancer cells depend on altered lipid metabolism for unrestrained growth and survival (48–50). However, how lipid metabolism is dysregulated and its functional significance in Ewing sarcoma remain poorly understood. In the present study, integrative analyses of LC-MS/MS-based lipidomics, RNA-Seq and ChIP-Seq together highlight that CRC factors converge on regulating key enzymes responsible for the biosynthesis of fatty-acids (FASN, SCD and ACLY), sphingolipids (SPTLC1, DEGS1 and UGCG) as well as glycerophospholipids (LPIN2). These lipid metabolism processes are required for cell survival of Ewing sarcoma, as evidenced by the cytotoxicity of their inhibitors (Orlistat against FASN and Myriocin against SPTLC1). In addition, silencing of either FASN, SCD or SPTLC1 markedly reduced cell proliferation of Ewing sarcoma, further corroborating this conclusion. The biochemical finding that KLF15 regulates lipid synthesis in Ewing sarcoma is not completely surprisingly given the established role of KLF15 in controlling lipid synthesis and fat storage in adipose tissues (51–54). However, to date, neither TCF4 nor NKX2-2 have been implicated in such metabolic processes in any cell type. These data together highlight CRC-lipid metabolism as a novel pro-tumor cascade in Ewing sarcoma.

As signature oncogenic pathways, PI3K/AKT and MAPK signaling pathways are important for almost every cancer type, including Ewing sarcoma (55–57). Our investigations (Fig. 7) uncover novel epigenomic mechanisms for the activation of these pathways by CRC factors in Ewing sarcoma. While TCF4 and NKX2-2 have not been involved in these pathways in any cell type, KLF15 is intriguingly reported to inhibit the AKT and MAPK signaling pathways in normal skeletal muscle, cardiomyocytes and kidney (58–60). It thus appears the regulatory effects of KLF15 on these signaling pathways are cell-type dependent, possibly due to the expression pattern of KLF15 target genes are cell-type specific. This is plausible given that KLF15 almost always cooperates with TCF4 and NKX2-2 in Ewing sarcoma in terms of genomic binding.

In conclusion, we identify and validate an Ewing sarcoma-specific CRC, which is under control of EWS-FLI1. Formed by KLF15, TCF4 and NKX2-2, this CRC apparatus coordinates the gene expression programs, including lipid metabolism, PI3K/AKT and MAPK signaling pathways, in Ewing sarcoma cells. These data advance the understanding of the mechanistic basis of transcriptional dysregulation in Ewing sarcoma, and provide potential novel therapeutic strategies against this malignancy.

## Materials and methods

### Cell culture

Ewing sarcoma cell lines (A673, SKNMC, EW8 and TC71) and human embryonic kidney cells 293T (HEK293T) used in this study were described previously (61, 62). Cells were grown in Dulbecco’s Modified Eagle Medium (DMEM) containing 10% fetal bovine serum (FBS), 100 U/ml penicillin and 100 mg/ml streptomycin, and kept in a humidified incubator at 37 °C with 5% CO_2_. All cell lines were recently authenticated by short tandem repeat analysis.

### Antibodies and reagents

The following antibodies were used in the current study: Anti-KLF15 (Proteintech, 66185-1-Ig), anti-TCF4 (Abcam, ab223073), anti-NKX2-2 (Proteintech, 13013-1-AP), anti-FLI-1 (Santa Cruz Biotechnology, sc-53826), anti-ACLY (Abcam, ab40793), anti-FASN (Abcam, ab22759), anti-ACC (Cell Signaling Technology, 4190S), anti-SPTLC1 (Proteintech, 66899-1-Ig), anti-SCD (Proteintech, 23393-1-AP), anti-FLAG (Sigma, F1804), anti-GAPDH (Cell Signaling Technology, 5174), anti-mouse IgG-HRP (Santa Cruz Biotechnology, sc-2005), anti-rabbit IgG-HRP (Santa Cruz Biotechnology, sc-2004) and rabbit IgG Isotype Control (Invitrogen, 02-6102), anti-mTOR (Cell Signaling Technology, 2972S), anti-P-mTOR (S2448) (Cell Signaling Technology, 2971S), anti-P-p38 MAPK (T180/Y182) (Cell Signaling Technology, 9216S), anti-p38 MAPK (Cell Signaling Technology, 8690S), anti-Akt (Cell Signaling Technology 4691S), anti-P-Akt (S473) (Cell Signaling Technology, 4060S), anti-p70 S6 Kinase (Cell Signaling Technology, 9202S) and anti-P-p70 S6 Kinase (Cell Signaling Techology, 9205S).

Reagents and kits included: Orlistat (Sigma-Aldrich, 96829-58-2), Myriocin, (Sigma-Aldrich, 35891-70-4), Propidium iodide (Sigma-Aldrich, 25535-16-4), FITC Annexin V Apoptosis Detection Kit (BD Biosciences), Dual-Luciferase Reporter Assay System (Promega), BioT transfection reagent (Bioland Scientific), Lipofectamine RNAiMAX transfection reagent (Invitrogen), and siRNA pools targeting *KLF15*, *TCF4*, *NKX2-2*, *FLI-1*, *FASN*, *SCD*, *RREB1* and *SPTLC1* (Dharmacon). siRNA sequences are provided in Supplementary Table 1.

### Construction of expression and lentiviral vectors

The pCDH-CMV-Flag-EF1-puro-KLF15 expression vector was amplified based on pCDH-CMV-Flag-EF1-puro vector, and a 3xFLAG-tag was added via PCR. pLKO.1-puro or pLKO-Tet-On vectors expressing shRNAs targeting *FLI1*, *KLF15*, *TCF4* and *NKX2-2* were constructed and confirmed by DNA sequencing. To produce viral particles, the recombinant viral vectors and packaging vectors were co-transfected into 293T cells. Supernatants containing viral particles were harvested at 48 hours and filtered through a 0.45 μM filter after transfection. A673 cells were then infected with the virus in the presence of 10 mg/ml polybrene.

### siRNA knockdown

Control scramble and target siRNAs were purchased from Shanghai Genechem Co. Cells were transfected with 100 nM siRNA in OPTIMEM-I (Gibco) for 24-48 hours using Lipofectamine RNAiMAX transfection reagent (Invitrogen). Two independent siRNA oligonucleotides were used for each gene.

### Chromatin immunoprecipitation (ChIP) and data analysis

ChIP was performed using our previously-described methods with slight modifications (11, 63). 2x10^7^-2x10^8^ of A673 cells were harvested and fixed with 1% paraformaldehyde for 10 min at room temperature. The fixation process was terminated by adding 250 mM of glycine. Chromatin solutions including lysis/wash buffer, sharing buffer and dilution buffer were prepared following a standard protocol (18). Samples were washed with PBS and lysed twice with lysis/wash buffer. After filtering through a 29 G needle, samples were harvested by spinning at 13,000 rpm for 10 min at 4°C. Sample pellets were then resuspended in sharing buffer and sonicated to shear genomic DNA to 300∼500bp. Sonicated sample lyses were subsequently spun at 13,000 rpm for 10 min in 4 °C to remove debris; the supernatants were diluted with dilution buffer. For immunoprecipitation, solubilized chromatin was incubated and rotated with 5 μg of anti-KLF15, anti-TCF4, anti-NKX2-2 antibody or IgG control antibody overnight at 4°C. Antibody-chromatin complexes were pulled down by Dynabeads Protein G (Life Technologies) for 4 hours at 4°C. The beads were washed with lysis/wash buffer followed by cold TE buffer. Finally, bound DNAs were eluted by elution buffer (1% SDS, 100 mM NaHCO_3_) and reverse-crosslinked overnight at 65 °C. DNA molecules were next treated with RNase A and proteinase K. Immunoprecipitated DNAs were extracted with the Min-Elute PCR purification kit (Qiagen), followed by either qPCR analysis, or DNA library preparation and sequencing on the HiSeq 4000 platform (Illumina).

ChIP-Seq was analyzed using established pipelines (11, 18). Raw reads were aligned to hg19 reference genome using bowtie 2 aligner (version 2.3.4.3) (64) followed by removal of PCR duplicates with Picard MarkDuplicates (http://broadinstitute.github.io/picard/). ChIP-Seq peaks were called using MACS (Model-Based Analysis of ChIP-Seq, version 2.1.2) (65) with default parameters. Reads were extended in the 5’ to 3’ direction by the estimated fragment length and normalized at -log 10 of the Poisson p-value of IP file compared to expected background counts by MACS2 bdgcmp command. BigWig files were generated by bedGraphToBigWig tool (https://genome.ucsc.edu/) and visualized in Integrative Genomics Viewer (http://www.broadinstitute.org/igv/home). Motif identification and comparison were performed with HOMER using findMotifsGenome program. H3K27ac ChIP-Seq data generated in 4 Ewing sarcoma cell lines, 3 primary tumors, primary mesenchymal stem cells (MSCs) and A673 cells were retrieved from NCBI Gene Expression Omnibus (GSE61953) and processed uniformly.

### Cell cycle analysis

Cells were harvested and washed with PBS, followed by fixation with 70% cold ethanol overnight at 4°C. Cells were washed twice with PBS and stained with propidium iodide. Cell cycle distribution was detected by SONY SA3800 spectral cell analyzer. Data were analyzed by FlowJo 7.6 software (Tree Star).

### Apoptosis assay

Cells were double stained with propidium iodide (PI) and Annexin V by using FITC Annexin V apoptosis detection kit (BD Biosciences) according to the manufacturer’s instructions. After staining, cells were analyzed using a BD FACSCanto II flow cytometer. Data were analyzed using FlowJo 7.6 software (Tree Star).

### RNA extraction and cDNA expression analysis

Total RNA was extracted using RNeasy mini kit (Qiagen). Purified RNA was reverse-transcribed using Maxima H Minus cDNA Synthesis Master Mix (Thermo Fisher). Quantitative real-time qPCR was performed on the AB7300 Detection System (Applied Biosystems, Foster City, CA) using gene-specific primers and Power SYBR Green PCR Master Mix (Applied Biosystems). Expression of each gene was normalized to *GAPDH*, and quantified using 2-delta (ct) method.

### Luciferase reporter assay

Candidate DNA regions (∼500bp) were PCR amplified and cloned into either pGL3-Promoter firefly luciferase reporter vector or pGL3-Basic luciferase reporter vector (Promega). Constructs were verified by Sanger sequencing. A673 cells were transfected using BioT transfection reagent. A Renilla luciferase control vector was co-transfected as a control for normalization. After 48 hours of transfection, the luciferase activity was measured by the Dual-Luciferase Reporter Assay System (Promega).

### Immunoblotting analysis

Whole cell lysates were prepared in RIPA buffer (1×PBS, 1% NP-40, 0.5% sodium deoxycholate, 0.1% SDS) supplemented with 10 mM beta-glycerophosphate, 1 mM sodium orthovanadate, 10 mM NaF, 1 mM phenylmethylsulfonyl fluoride (PMSF), and 1×Roche Protease Inhibitor Cocktail (Roche, Indianapolis, IN). Immunoblotting was performed using specific primary antibodies as indicated and horseradish peroxidase (HRP)-conjugated secondary antibodies.

### Xenograft assays in nude mice

All animal experiments were conducted following protocols approved by the Institutional Animal Care and Use Committee (IACUC) of Cedars-Sinai Medical Center. For the study of Orlistat treatment in A673 xenografts, twelve 5-6 weeks old BALB/c-nu female mice (Taconic Bioscience) were subcutaneously inoculated in their dorsal flanks with a suspension of A673 cells (2.0×10^6^). When the tumors grew to 50 mm^3^, mice were injected intraperitoneally with either Orlistat (200 mg/kg/day) or equal volume of vehicle (10% DMSO, 20% cremophor and 70% NaCl). For the study of either KLF15 or TCF4 knockdown, A673 cells were engineered to stably express either doxycycline (DOX)-inducible scrambled shRNA or shRNAs against either *KLF15* or *TCF4*, and were cultured in DMEM supplemented with 10% Tetracycline (Tet)-free FBS (Biological Industries) before implantation. After tumor inoculation, mice were randomized into two groups, and were fed with 2.5 mg/ml doxycycline containing water to turn on the expression of shRNAs. Tumor size and body weight were measured every 2 days. Mice were sacrificed by CO_2_ inhalation when the largest tumors were approximately 1.5 cm in diameter; tumors were dissected, weighted, and analyzed.

### RNA-Seq and data analysis

RNAs were isolated using miRNeasy RNA isolation kit (Qiagen). We aligned 150bp paired-end reads to hg19 (UCSC) genome using Kallisto psedo aligner (66). Reads were counted with tximport Bioconductor package (67) and normalized to gene levels by Transcript level abundances (TPM). Differentially expressed genes were identified by DESeq2 package (68) with adjusted p value < 0.05 and absolute log 2 (fold change) > 0.5. Pathway enrichment analysis was performed using ConsensusPathDB (http://cpdb.molgen.mpg.de/) and KEGG. For GSEA analysis, we used significantly downregulated genes (adjusted p value < 0.05 and log 2 (fold change) < -1) in each knockdown of TF as the annotation base and performed enrichment analyses in all expressed genes with mean TPM values > 0.5 in other two TFs.

### Computational construction of CRC

CRC was computationally constructed using our published methodology (18, 24, 25) with important modifications. Given the central role of EWS-FLI1 in establishing the transcriptome of Ewing Sarcoma cells, we required that all candidate CRC members have both EWS-FLI1 binding motif and binding peaks in their super-enhancer regions. Therefore, we focused on A673 cell line, a well-characterized Ewing Sarcoma line which had matched H3K27ac and EWS-FLI1 ChIP-Seq results (7, 69). Briefly, we first identified 77 super-enhancer-assigned TFs in A673 cell line as shown in Supplementary Table 2. Next, super-enhancer regions assigned to these TFs were extended 500 bp both upstream and downstream, followed by motif scanning with FIMO, for the identification of super-enhancer-assigned auto-regulated TFs. Finally, motif scanning was applied to identify further potential binding sites from other TFs of all auto-regulated TFs in their extended super-enhancer regions which, as mentioned above, must also have EWS-FLI1 binding motif and peaks. Circuitries were then constructed based on all possible fully interconnected auto-regulatory loops.

### Hi-C interactions

The Hi-C data of SKNMC cell line were downloaded from ENCODE database and processed based on the pipeline published by Dixon et al (69). We extracted the interactions of the loci of *KLF15*, *TCF4* and *NKX2-2* at 40 kb revolution and visualized in UCSC genome Browser (https://genome.ucsc.edu/index.html).

### Liquid chromatography tandem mass spectrometry (LC-MS/MS)-based lipidomics

Lipidomic analysis was performed as described previously with slight modifications (70). Briefly, total cellular lipids were extracted with methyl tert-butyl ether (MTBE) (Sigma Aldrich) from fresh cell pellets and dried in a SpeedVac concentrator (Thermo Scientific). Lipid samples were resuspended in 50% isopropanol, 50% methanol and analyzed by liquid chromatography tandem mass spectrometry (LC-MS/MS). Twenty microliters of lipid solution were loaded onto a 15 cm Accucore Vanquish C18 column (1.5 μm particle size, 2.1 mm diameter) and separated using an Ultimate 3000 XRS ultraperformance LC system (Thermo Scientific). The mobile phase consisted of 60% acetonitrile, 10 mM ammonium formate, and 0.1% formic acid (phase A) and 90% isopropanol, 10% acetonitrile, 10 mM ammonium formate, and 0.1% formic acid (phase B). LC gradient was 35-60% B for 4 min, 60-85% B for 8 min, 85%-100% for 9 min, 100% B for 3 min, 100-35% B for 0.1 min, and 35% B for 4 min at a flow rate of 0.3 ml/min. Mass spectra were acquired by an Orbitrap Fusion Lumos Tribrid mass spectrometer (Thermo Scientific) operated in a data-dependent manner. Parameter settings for FTMS1 included orbitrap resolution (120,000), scan range (m/z 250-1200), AGC (2×10^5^), maximum injection time (50 ms), RF lens (50%), data type (profile), dynamic exclusion for 8s using a mass tolerance of 25 ppm, and cycle time (2 s); FTMS2 included orbitrap resolution (30,000), isolation window (1.2 m/z), activation type (HCD), collision energy (30±3%), maximum injection time (70 ms), AGC (5×10^4^), and data type (profile). Acquired raw files were analyzed using LipidSearch (v1.4) (Thermo Scientific) for sample alignment, MS2 identification, and MS1 peak area calculation. Statistical analyses were conducted using the Perseus (v1.6.6.0) software (71), wherein the p values were calculated by two-tailed Student’s t-test and corrected for multiple hypothesis testing via the Benjaminin-Hochberg method. Volcano plots were generated using the ggplot2 in the R environment (R Development Core Team; https://www.r-project.org/) (v3.5.0).

### Statistical analysis

Two-tailed Student’s t-test was used to evaluate the statistical difference between two groups, while one-way analysis of variance (ANOVA) was applied for multi-group comparisons. All statistical analyses were performed with SPSS 19.0. Log-rank test was used for survival analysis. Differences were considered statistically significant at p < 0.05 (*), p < 0.01 (**) and p < 0.001 (***); n.s., not significant. Diagrams were created by GraphPad Prism 6.

## Data availability

ChIP-Seq data of KLF15, TCF4 and NKX2-2, and RNA-Seq data of A673 upon knockdown of each TF have been deposited into the GEO under accession number GSE141493. For ChIP-Seq data, the matching input file was obtained from our previous published data (GSM2944109).

## Author contributions

D.-C.L. conceived and devised the study. D.-C.L., X.P.S. and Y.Y.Z. designed experiments and analysis. X.P.S. and L.L.J., performed the experiments. W.Y. and B.Z. performed quantitative lipidomic and data analysis. Y.Y.Z. and M.L.H. performed bioinformatics and statistical analysis. S.G. contributed reagents and materials. X.P.S., Y.Y.Z., L.L.J., D.-C.L., and H.P.K. analyzed the data. D.-C.L. X.P.S. and H.P.K. supervised the research and wrote the manuscript.

## Acknowledgements

This research is supported by the National Research Foundation Singapore under its Singapore Translational Research (STaR) Investigator Award (NMRC/STaR/0021/2014) and administered by the Singapore Ministry of Health’s National Medical Research Council (NMRC), the NMRC Centre Grant awarded to National University Cancer Institute of Singapore, the National Research Foundation Singapore and the Singapore Ministry of Education under its Research Centres of Excellence initiatives (to H.P.K). This work was also supported by NIH grant (1R01 CA200992) to H.P.K. This work is additionally supported by NSFC (81670154/H0812,81470355/H1616), Projects (201707010352) from the Foundation of Guangzhou Science and Technology Innovation Committee in China (to X.S.). This research was also supported by Alan B. Slifka Foundation and the Ewing’s Sarcoma Research Foundation.

