## Supplemental figure and figure legends for "EWS-FLI1 regulates and cooperates with core regulatory circuitry in Ewing sarcoma"

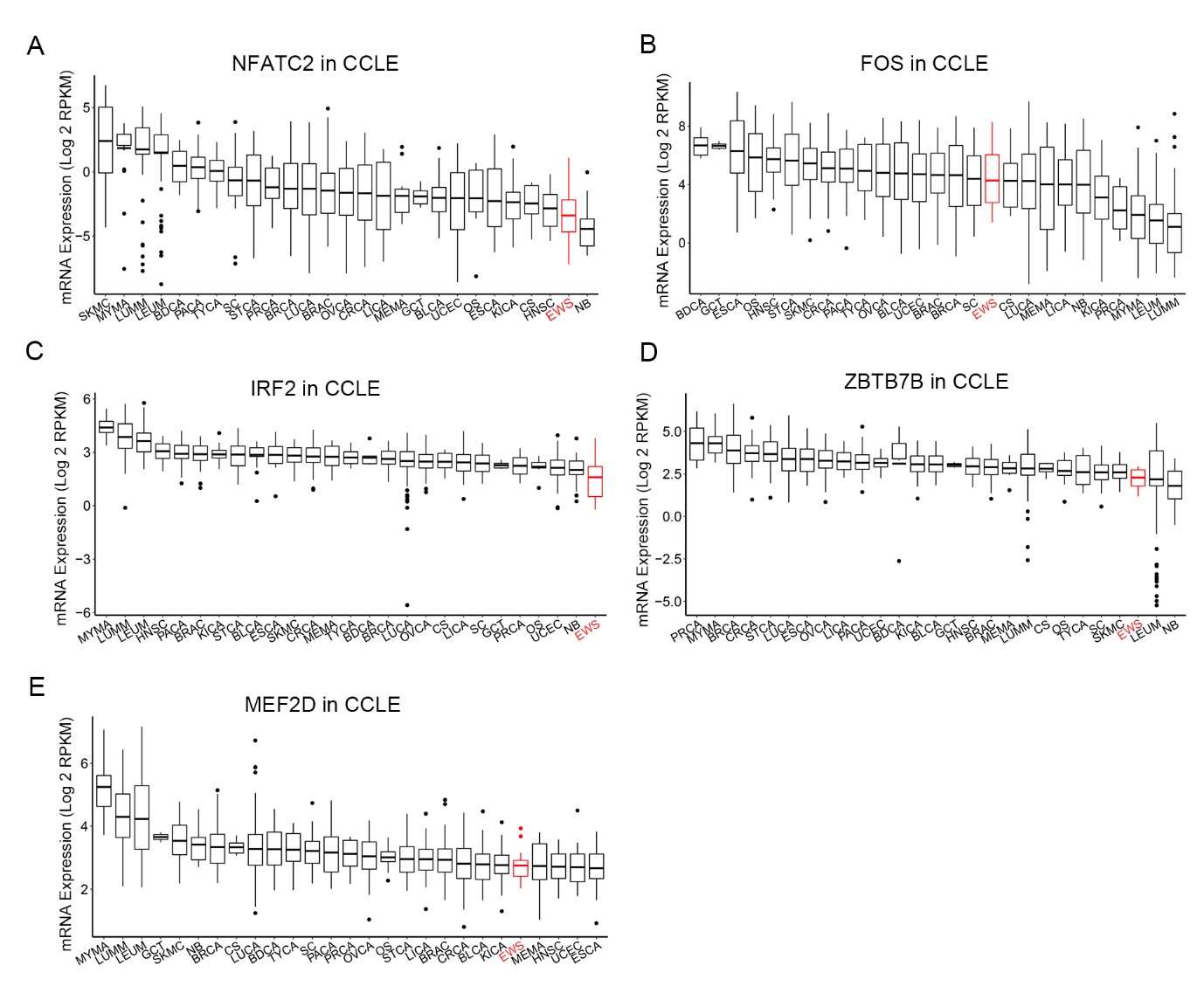

**Supplement Figure 1.** The mRNA levels of candidate TFs (NFATC2, FOS, IRF2, ZBTB7B and MEF2D) in CCLE database.

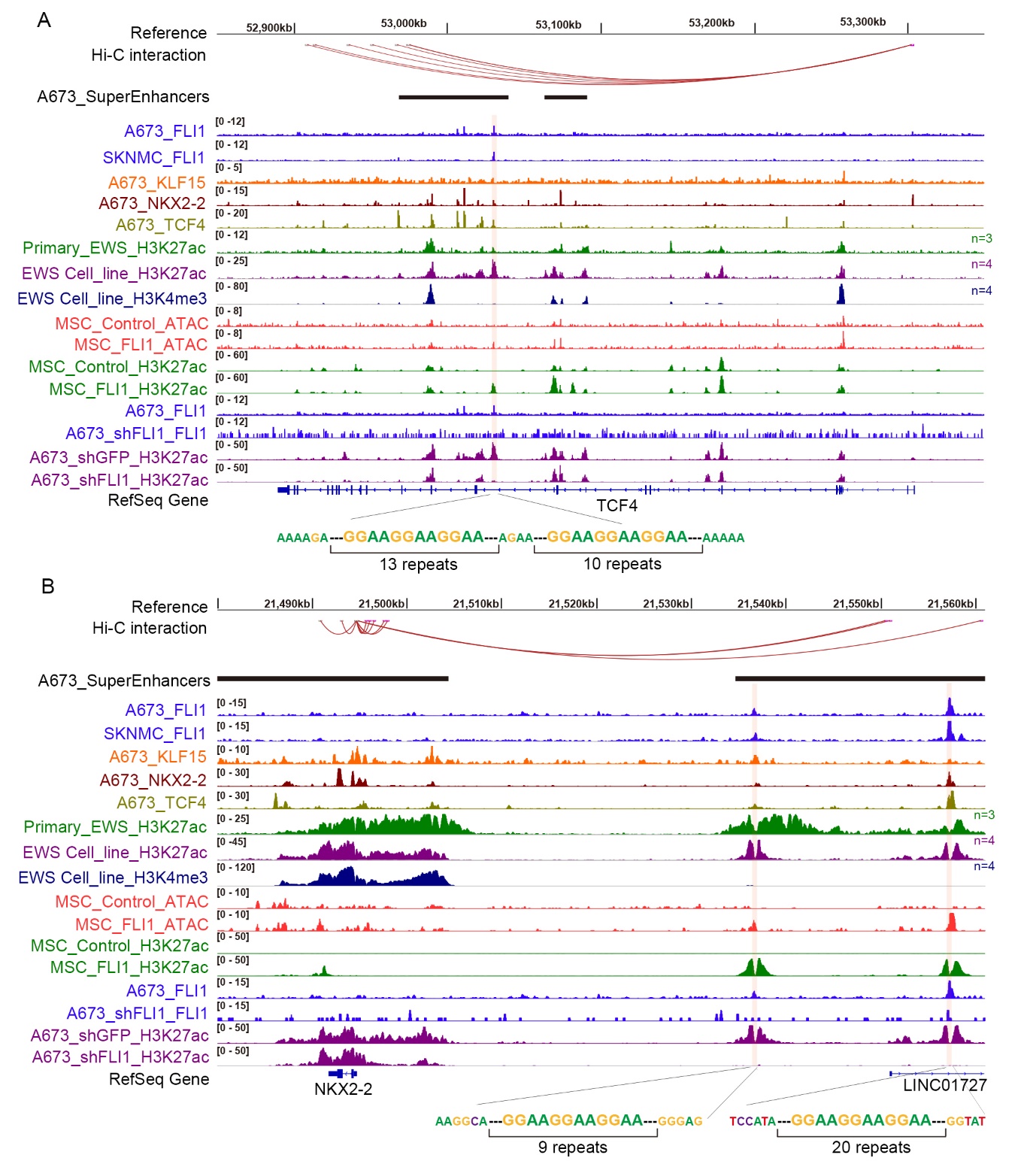

**Supplement Figure 2**. **Co-binding peaks of** **EWS-FLI1 and CRC TFs at the genomic loci of TCF4 and NKX2-2**. IGV plots of ChIP-Seq showing co-occupancy of EWS-FLI1 and CRC TFs at the super-enhancers and promoters of TCF4 (A) and NKX2-2 (B). Hi-C interactions were re-analyzed from the data of SKNMC cell line downloaded from ENCODE database; H3K27ac, H3K4me3 and EWS-FLI1 ChIP-Seq data were retrieved from GEO (GSE61953). ATAC-Seq and ChIP-Seq profiles at super-enhancers of TCF4 and NKX2-2 in the presence and absence of either EWS-FLI1 overexpression or knockdown. Data were retrieved from GEO (GSE61953).

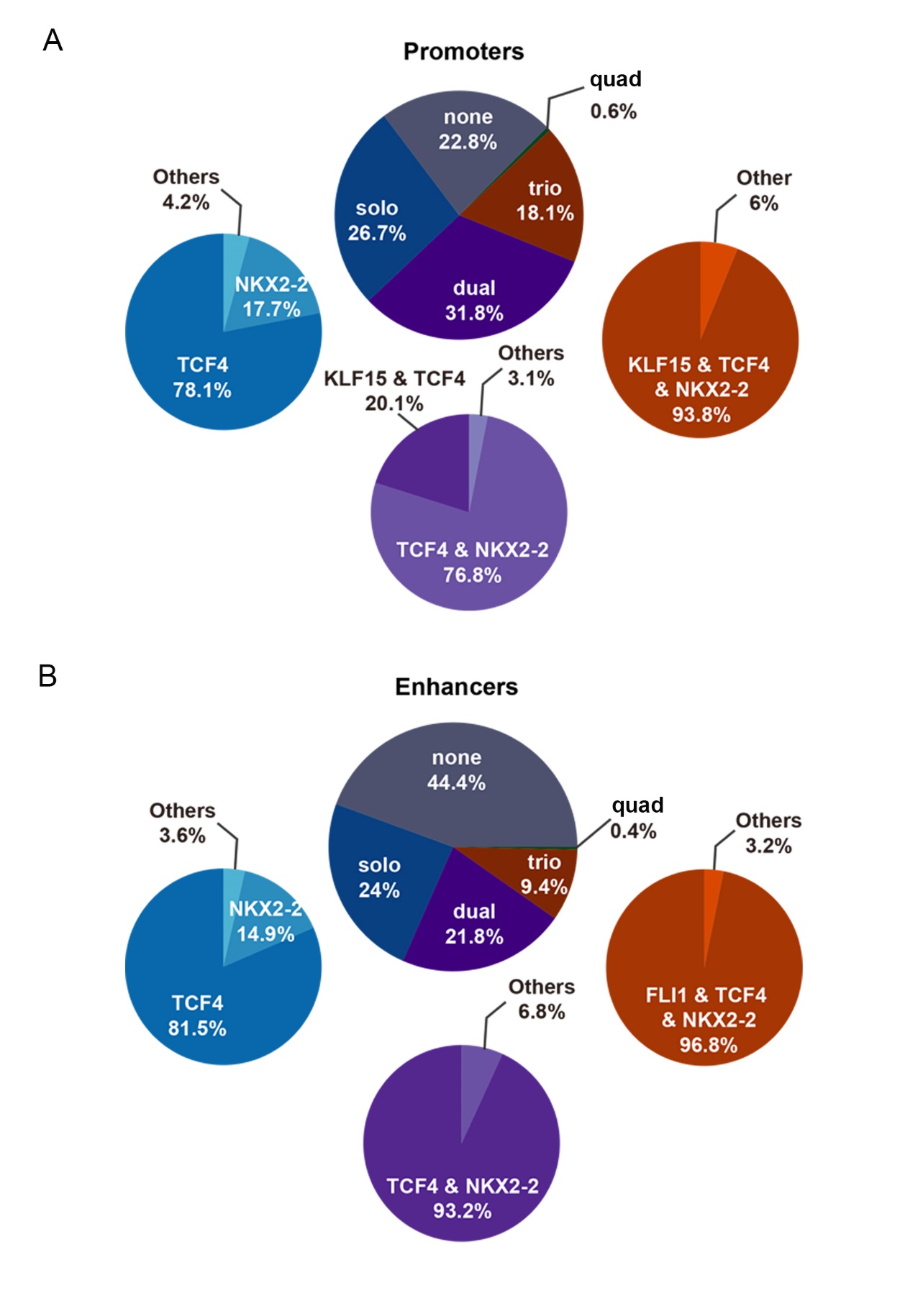

**Supplement Figure 3.** Pie charts of the factions of combinatorial binding patterns of EWS-FLI1 and three CRC TFs in both enhancers and promoters.

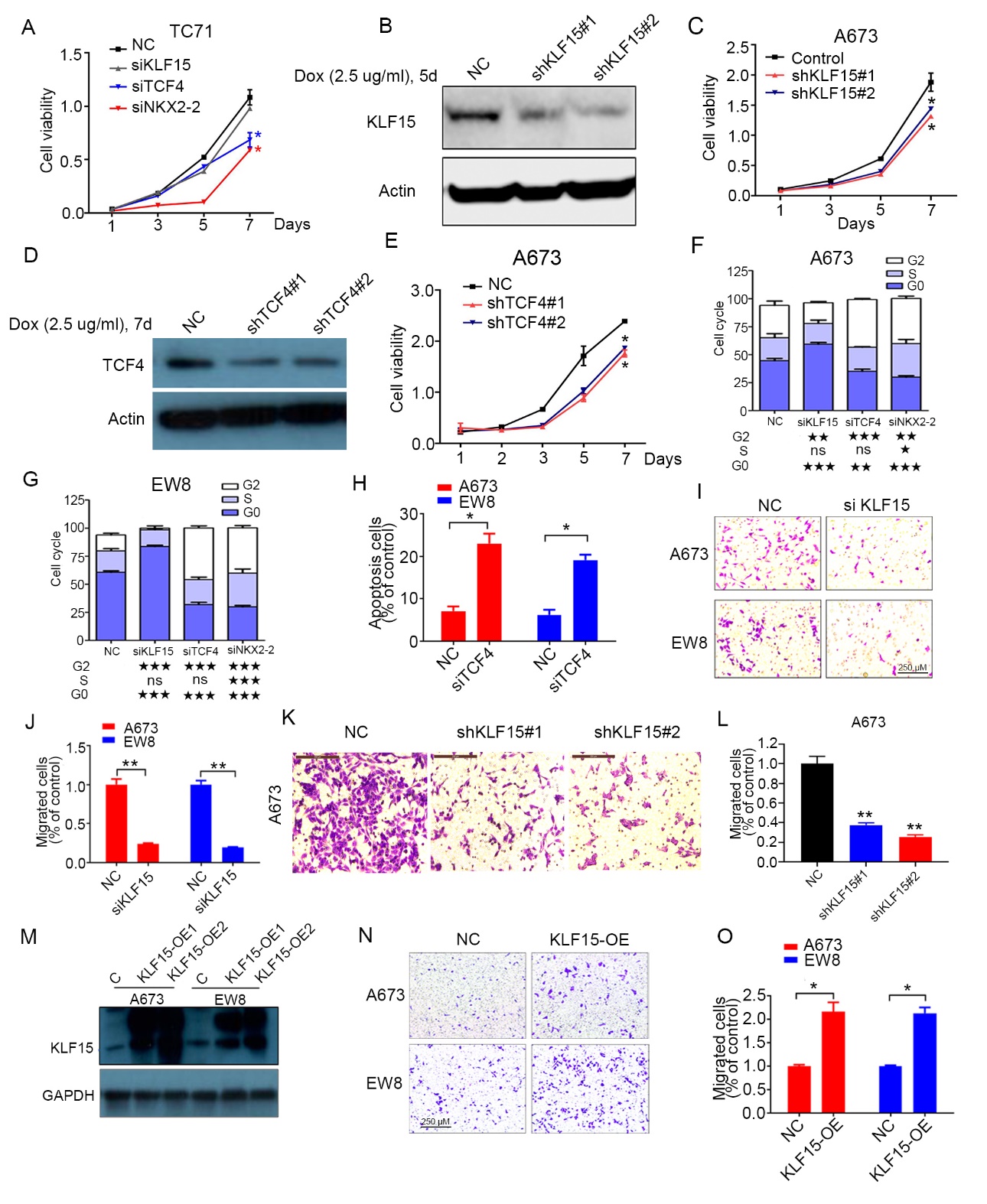

**Supplement Figure 4. The cancer-promoting functions of KLF15 and TCF4 in Ewing sarcoma cells.** (A) Silencing of each of three CRC TFs by siRNAs decreased cell proliferation in TC71 cells. (B-E) Knockdown of KLF15 (B, C) or TCF4 (D, E) by inducible shRNAs decreased cell proliferation in A673 cells. (F, G) Knockdown of three CRC TFs by individual siRNAs increased cell-cycle arrest in Ewing sarcoma cell lines. (H) Silencing of TCF4 by siRNAs induced cell apoptosis in A673 and EW8 cells. Cells were stained with Annexin V/PI. Mean ± s.d. are shown, n = 6. *, P < 0.05; **, P < 0.01. (I-L) Silencing of KLF15 by siRNAs (I, J) or inducible shRNAs (K, L) decreased cell migration as measured by transwell assays. (M-O) Forced expression of KLF15 enhanced cell migration in A673 cells. Mean ± s.d. are shown, n = 3. *, P < 0.05; **, P < 0.01.

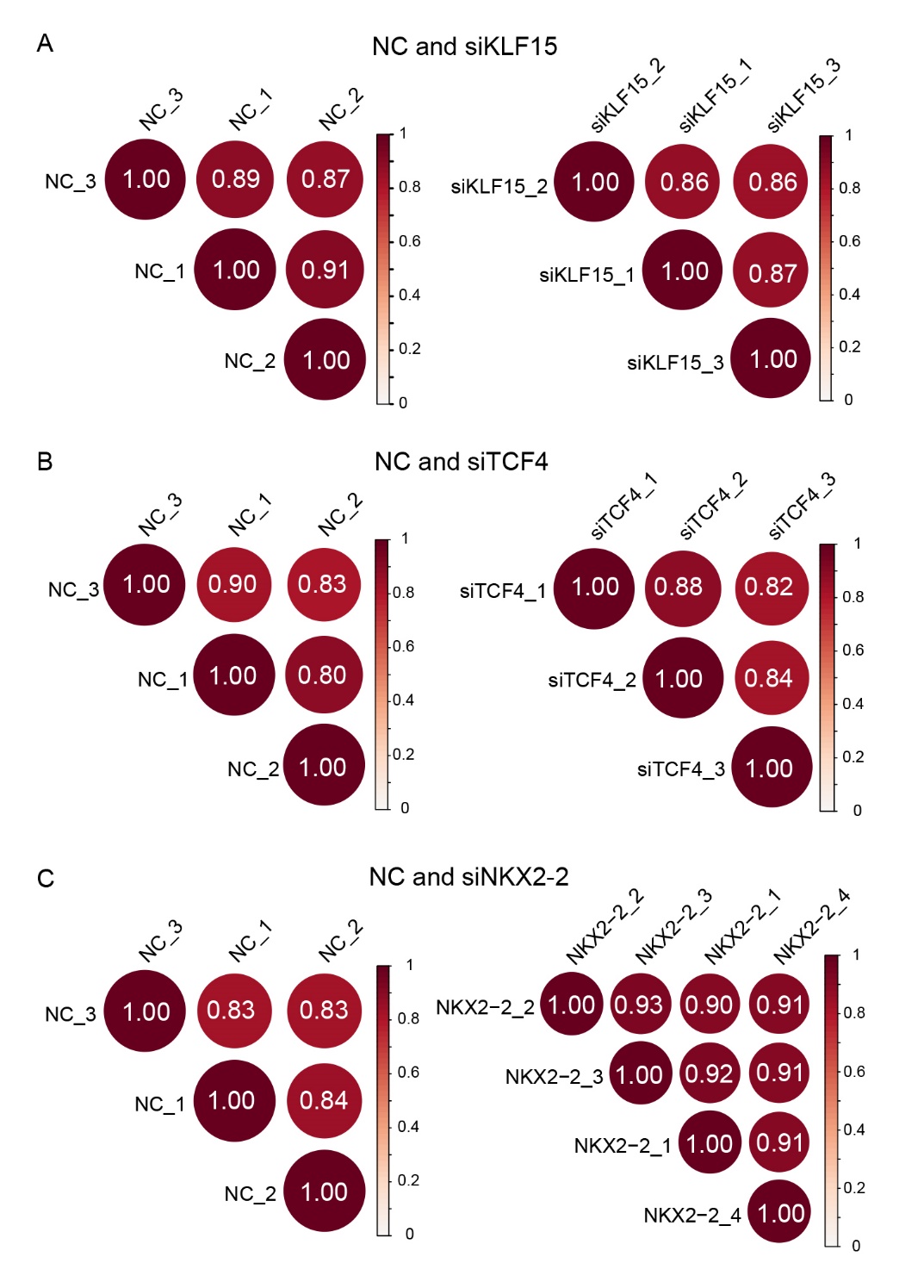

**Supplement Figure 5**. Pearson correlation plots between replicates of LC-MS/MS-based lipidomic samples from each group.
