## Supplemental Table1 for "EWS-FLI1 regulates and cooperates with core regulatory circuitry in Ewing sarcoma"

### Supplementary Table 1: siRNA sequences

| Target | sense（5'-3'） | antisense（5'-3'） |
|---|---|---|
| siFLI1 #1 | GGGAGUAUGACCACAUGAATT | UUCAUGUGGUCAUACUCCCTT |
| siFLI1 #2 | GCCAUAAAGGAGUACAGCUTT | AGCUGUACUCCUUUAUGGCTT |
| siKLF15 #1 | GCAUCUUGGACUUCCUAUUTT | AAUAGGAAGUCCAAGAUGCTT |
| siKLF15 #2 | GCUUGCCCGAGUUUCCUUUTT | AAAGGAAACUCGGGCAAGCTT |
| siTCF4 #1 | GGGACAUGCAUGGAAUCAUTT | AUGAUUCCAUGCAUGUCCCTT |
| siTCF4 #2 | CUCAUCGUCUCCUAAUUAUTT | AUAAUUAGGAGACGAUGAGTT |
| siNKX2-2 #1 | CCUGCCGGACACCAACGAUTT | AUCGUUGGUGUCCGGCAGGTT |
| siNKX2-2 #2 | GCACCGAGGGCCUUCAGUATT | UACUGAAGGCCCUCGGUGCTT |
| siSCD #1 | GACGAUAUCUCUAGCUCCUTT | AGGAGCUAGAGAUAUCGUCTT |
| siSCD #2 | GGUUGAAUAUGUCUGGAGATT | UCUCCAGACAUAUUCAACCTT |
| siFASN #1 | GGACCUGUCUAGGUUUGAUTT | AUCAAACCUAGACAGGUCCTT |
| siFASN #2 | CCCAGGCUGAAGUUUACAATT | UUGUAAACUUCAGCCUGGGTT |
| siRREB1 #1 | GAGCGAACCUUCACCUUGATT | UCAAGGUGAAGGUUCGCUCTT |
| siRREB1 #2 | GACCUAUCUUCCAUCAACATT | UGUUGAUGGAAGAUAGGUCTT |
| siSPTLC1 #1 | GAUCUGAUCUUACAGUCAATT | UUGACUGUAAGAUCAGAUCTT |
| siSPTLC1 #2 | GGAUUGUUGGAUAACCCUATT | UAGGGUUAUCCAACAAUCCTT |
